# Deciphering the Genomic Landscape of Renal Cell Carcinoma Brain Metastases

**DOI:** 10.64898/2026.05.02.722447

**Authors:** Betul Gok Yavuz, Peng Li, Jose A Ovando-Ricardez, Alessandro La Ferlita, Jonathan W. T. Tse, Sahin Hanalioglu, Berrin Babaoglu, Baylar Baylarov, Lisa M. Norberg, Haidee D. Chancoco, Erika J. Thompson, Melike Mut, Figen Soylemezoglu, Jason T. Huse, Adeboye O. Osunkoya, Mehmet Asim Bilen, Merve Hasanov, Eric Jonasch, David J. H. Shih, Elshad Hasanov

**Affiliations:** Division of Medical Oncology, Department of Internal Medicine, The Ohio State University Comprehensive Cancer Center, Columbus, OH, USA; Pelotonia Institute for Immuno-Oncology, The Ohio State University Comprehensive Cancer Center, Columbus, OH, USA; Section of Hematology/Oncology, Department of Medicine, University of Chicago, Chicago, IL, USA; School of Biomedical Sciences, Li Ka Shing Faculty of Medicine, The University of Hong Kong, Hong Kong SAR, China; Department of Neurosurgery, Hacettepe University Faculty of Medicine, Ankara, Turkey; Department of Pathology, Hacettepe University Faculty of Medicine, Ankara, Turkey; Department of Pathology, The University of Texas MD Anderson Cancer Center, Houston, TX, USA; Cancer Neuroscience Program, The University of Texas MD Anderson Cancer Center, Houston, TX, USA; Department of Epidemiology, The University of Texas MD Anderson Cancer Center, Houston, TX, USA; Department of Genetics, The University of Texas MD Anderson Cancer Center, Houston, TX, USA; Department of Neurosurgery, The University of Virginia Health System, Charlottesville, VA, USA; Departments of Pathology and Urology, Emory University School of Medicine, Atlanta, GA, USA; Department of Hematology and Medical Oncology, Emory University School of Medicine, Atlanta, GA, USA; Department of Genitourinary Medical Oncology, Division of Cancer Medicine, The University of Texas MD Anderson Cancer Center, Houston, TX, USA

**Keywords:** renal cell carcinoma, brain metastases, whole-exome sequencing, copy number alterations, cancer genomics

## Abstract

Brain metastases from renal cell carcinoma (RCC) remain a major cause of morbidity and mortality, yet the genomic features associated with metastatic dissemination remain poorly understood. Whole-exome sequencing was performed on 72 RCC brain metastasis samples with matched normal. To identify candidate genomic alterations associated with brain metastasis, the genomic alterations detected in the brain metastases were compared against alterations in extracranial metastases from the MSK-ECM cohort (n=137) and primary RCC tumors from TCGA (n=432) by case-control analyses. Candidate alterations were also identified through matched-pair analyses comparing brain metastases with matched primary tumors or extracranial metastases from the same patient (n=25). A random survival forest model incorporating the candidate CNA events was developed to predict overall survival. The candidate CNAs were further evaluated using functional experimental data from MetMap and DepMap. Survival analyses were conducted to assess the prognostic relevance of these alterations. We identified recurrent CNAs enriched in RCC brain metastases, including 4q loss, 7p gain, 7q gain, 8p loss, 8q gain, 9p21.3 deletion, 12q15 amplification, and 14q loss. These alterations were associated with significantly poorer patient survival among RCC patients. A random survival forest model based on these CNA events stratified TCGA-KIRC patients into prognostically distinct risk groups (C-index = 0.64). Among the recurrent CNAs, 8p loss, 8q gain, 9p21.3 deletion were associated with increased incidence of brain metastases across multiple primary cancer types in xenograft mouse models. These alterations were also strongly associated with metastatic progression and poor prognosis across RCC, lung adenocarcinoma, breast cancer, and melanoma. These findings indicate a shared genomic basis for brain tropism and highlight the potential utility of copy-number alterations as biomarkers for risk stratification and clinical decision-making.

## INTRODUCTION

Brain metastases from renal cell carcinoma (RCC) cause substantial morbidity and remain a therapeutic challenge ^1,2^. Historically reported in 2–15% of patients ^1^, recent studies indicate that brain metastases now occur in 20–28% of RCC cases ^3,4^, underscoring their rising clinical relevance. Prognoses for brain metastasis patients remain poor. In the JAVELIN Renal 101 trial, patients with asymptomatic brain metastases had markedly shorter progression-free survival (PFS) at 2.8–4.9 months, compared with 8.4–13.8 months in the overall trial population ^5^.

Limited treatment efficacy against RCC brain metastases stems from their biological complexity and tumour heterogeneity. Although targeted therapies and immunotherapies have transformed primary RCC management, their efficacies against brain metastases remain modest ^1,6,7^, leaving radiation and surgery as the mainstays of care. This underscores the need to understand the genomic landscape of RCC brain metastasis, which can differ substantially from primary tumors and extracranial metastases ^8,9^.

The TRACERx study identified 9p and 14q loss as key drivers of RCC metastasis and poor prognosis ^9,10^; however, it included only one patient with brain metastasis, leaving the genomic basis of CNS spread largely unexplored. To address this gap, we performed whole-exome sequencing (WES) of RCC brain metastases, primary tumors, and extracranial metastases to identify copy number alterations (CNAs) associated with brain metastasis, which may guide future development of therapeutic strategies.

## METHODS

Detailed methodology is provided in the Supplementary Methods.

### Study cohort

Tissue specimens were collected under individual Institutional Review Board (IRB) approved protocols at the University of Texas – MD Anderson Cancer Center, Emory University and Hacettepe University, and all processing procedures performed on patient samples were in accordance with the ethical standards of the Helsinki Declaration and its later amendments. Consent was obtained, and annotation of specimens was performed via standard operating procedures of the respective Biobank framework and personnel, with the consent obtained through either in-person signatures or IRB-approved remote consent procedures. The study cohort is explained in Figure 1A. Histologic subtypes of samples are shown in **Supplementary Table 1**.

**Figure 1.**
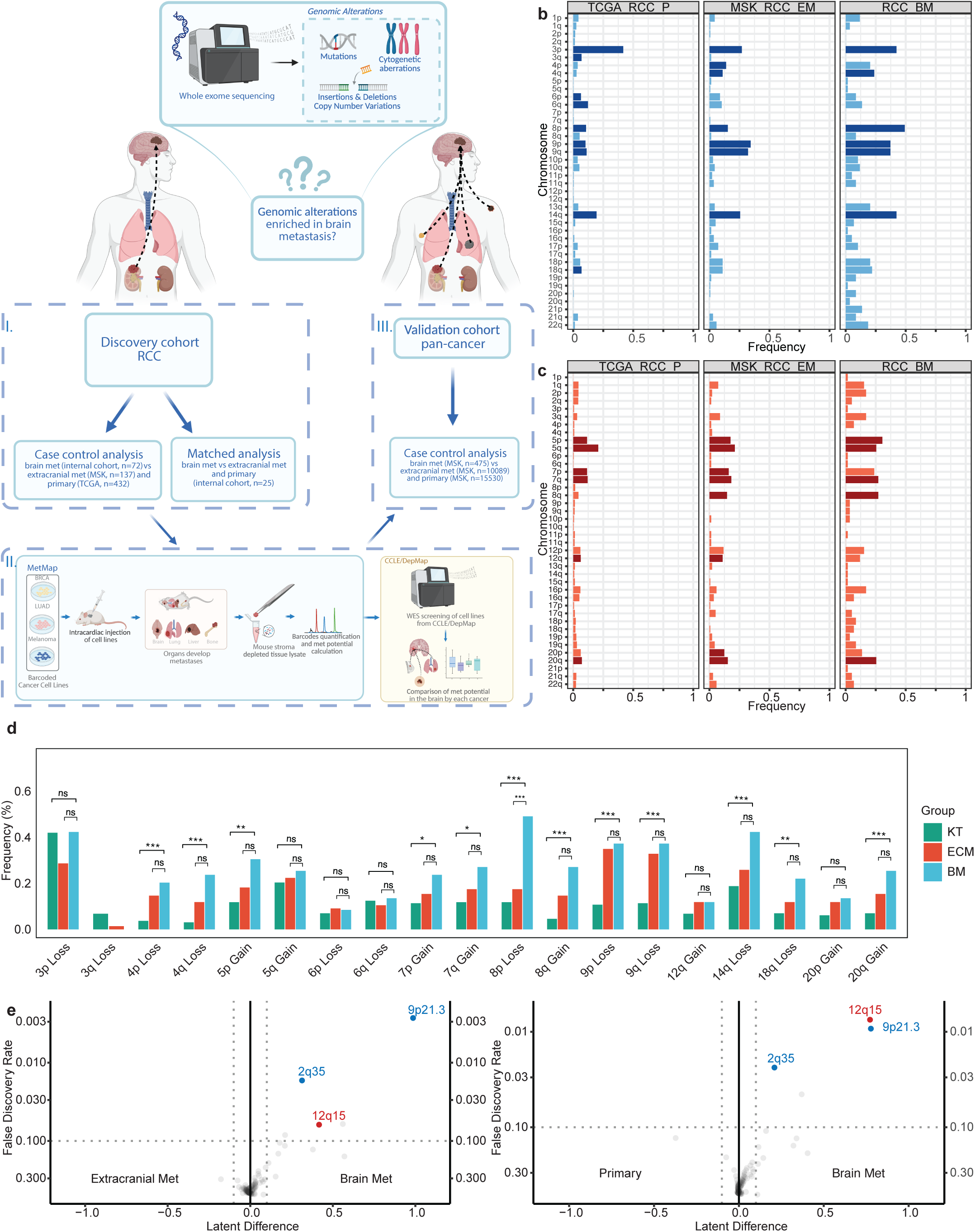
**A**. Schematic overview of the study design and workflow to identify genomic alterations driving brain metastases in renal cell carcinoma (RCC). The study utilized whole-exome sequencing (WES) to detect single nucleotide variations (SNVs) and copy number alterations (CNAs). The study was conducted in three phases: (I) A discovery cohort of RCC brain metastases (BM) was analyzed through case-control comparisons of BM samples with extracranial metastases from MSK dataset and primary RCC tumors from TCGA. Matched analysis was performed on 25 brain metastasis samples paired with primary and/or ECM tumors to identify unique genomic events. (II) Findings from the discovery cohort were further tested in other cancer types using MetMap analysis. (III) Findings from discovery cohort and MetMap analysis were validated in a pan-cancer cohort using MSK datasets. The Frequency of chromosomal arm level losses (**B**) and gains (**C**) in RCC-BM, TCGA-KIRC and MSK-ECM. Significant cytogenetic alterations (bold colors) were identified using the exact binomial test by comparing the frequency of cytogenetic alteration against the background frequency estimated based on gene content (FDR< 10%). Comparison of proportion of significant chromosomal arm level CNAs across these three cohorts was done by using Chi Square test (**D**). Significant focal CNAs in RCC-BM vs TCGA-KIRC and BM vs MSKCC-ECM (red showing gains, blue showing losses) (**E**). *p<0.05, **p<0.001, ***p<0.0001.

### Sample preparation and sequencing

DNA was extracted from frozen and FFPE tumor tissue using commercial kits following institutional biobank protocols, requiring tumor purity ≥40%. Matched normal tissue was confirmed by genotyping. Whole-exome libraries were prepared with Twist library preparation and exome capture kits, followed by Illumina NovaSeq 6000 paired-end sequencing. Library quality, concentration, and target enrichment were assessed using fluorometric assays, TapeStation, and qPCR. Full sequencing procedures are provided in the Supplementary Methods.

### Copy-number and single-nucleotide variant analysis

Copy-number analysis of WES data followed modified versions of established pipelines ^11^. Read coverage across exome targets (±250 bp) was obtained using GATK4-CNV and normalized to diploid controls to generate total copy-ratio profiles, which were segmented using circular binary segmentation ^12^. Allele-specific copy numbers were inferred by combining total copy ratios with B-allele frequencies from germline heterozygous SNPs ^11^, and ASCAT was used to generate allele-specific CNA profiles; rescaled copy-number values were computed as previously described ^11^.

Segmented copy-number and gene-level profiles for TCGA and MSK cohorts were retrieved from PanCanAtlas ^13^ and cBioPortal ^14^. Arm-level cytogenetic alterations were defined using segment-length–weighted mean copy-number states ^15^. For focal loci (2q35, 9p21.3, 12q15), gene identities were extracted using *Organism.dplyr*, and locus-level copy number was assigned by averaging gene-level values from cBioPortal^14^.

Somatic SNVs were analyzed using *dNdScv* ^16^ to identify genes mutated at frequencies exceeding background.

### Co-occurrence analysis

To characterize genomic co-alteration patterns, we constructed a binary event matrix for recurrent arm-level and focal CNAs in RCC brain metastases. Pairwise co-occurrence was evaluated using a probabilistic framework, and significant associations were visualized as a network. These patterns informed downstream classification of RCC-BM samples into CNA-defined subgroups for oncoprint visualization.

### Comparison of mutation frequencies in the PI3K/AKT/mTOR pathway

We compared the mutation frequencies of MTOR, PIK3CA, and PTEN between our RCC brain metastasis cohort and primary RCC tumors from the TCGA-KIRC cohort. Mutation data for TCGA-KIRC were retrieved from cBioPortal.

### CNA-based survival modeling using Random Survival Forests

A random survival forest (RSF) model incorporating these CNAs was trained and validated following established protocols for genomic survival modeling, with performance summarized using the concordance index. Detailed statistical specifications, model parameters, and multiple-testing procedures are provided in the Supplementary Methods.

### MetMap

To evaluate whether candidate CNAs associated with RCC brain metastasis generalize across other cancers, we integrated organ-specific metastatic potential scores from the MetMap500 project ^17^ with corresponding copy-number profiles from DepMap. LUAD, BRCA, and melanoma cell lines were stratified by the presence of 8p loss, 8q gain, or 9p21.3 deletion, and brain, liver, lung, and bone metastatic-potential scores were compared using Wilcoxon rank-sum tests.

### Statistical analysis

All analyses were performed in R (v3.2.3). Two-sided tests were used throughout, and P < 0.05 was considered statistically significant unless otherwise specified. CNA and mutation enrichment across cohorts was assessed using weighted logistic regression with matching weights, or Wilcoxon rank-sum tests for continuous metastatic-potential scores.

For survival analysis, OS and PFI metrics were obtained from TCGA-KIRC and MSK cohorts. Patients lacking all candidate events (4q loss, 7p gain, 7q gain, 8p loss, 8q gain, 9p21.3 deletion, 12q15 gain, 14q loss) served as the reference group. Associations between CNAs and survival were assessed using Kaplan–Meier estimates, log-rank tests, and Cox-based hazard ratios. Details are provided in the Supplementary Methods.

## RESULTS

### Case-control analyses identified copy-number alterations enriched in brain metastases in RCC

To identify genomic alterations that drive RCC-brain metastasis, we performed WES in 72 brain metastasis samples (**Fig. 1A**), comprehensively characterizing focal and broad copy number alterations, as well as single nucleotide variations (**Supplementary Fig. 1A-B**). We conducted this study in three phases. First, we constructed a discovery cohort consisting three groups: RCC brain metastases (RCC-BM; n=72), extracranial metastases from MSKCC (MKS-RCC-EM; n=137), and primary RCC tumors from TCGA (TCGA-RCC-P; n=432) ^18,19^. We performed case-control analyses comparing brain metastases against extracranial metastases and primary tumors. We also performed matched-pair analyses on 25 brain metastases with matched primary or extracranial samples. Second, candidate genomic alterations associated with brain metastasis identified in the discovery cohort were evaluated using MetMap, which interrogated the propensity of genomically characterized cells lines to metastasize to different organs in xenograft mouse experiments ^20,21^. Third, the candidate genomic alterations were validated in a non-overlapping pan-cancer cohort of brain metastasis (n=475) extracted from the MSKCC dataset.

We first identified recurrent copy-number alterations (CNAs) that occur at significantly higher frequencies vs. background, as previously described ^15^ (**Fig. 1B-C**). This analysis confirmed the cytogenetic hallmarks of RCC: 3p loss and 5q gain, as recurrent CNAs in all three groups (TCGA-RCC-P, MSK-RCC-EM, and RCC-BM) in the discovery cohort.

Owing to positive selection during metastatic dissemination to the brain, we expect metastatic drivers to occur across brain metastases at significantly elevated frequencies compared against extracranial metastases and primary tumours (**Supplementary Fig. 1C-D**). Therefore, we compared the frequencies of recurrent CNAs across these three cohorts and identified CNAs that were enriched in brain metastases, including losses of 4p, 4q, 8p, 9p, 9q, 14q and 18q, as well as gains of 5p, 7p, 7q, 8q, and 20q (**Fig. 1D**). Notably, 8p loss was the most frequent BM CNA event, observed in ∼50% of brain metastasis samples vs. 15% in primary and ECM samples (**Fig. 1C-D**).

Furthermore, we identified distinct focal regions with significantly higher positive selection scores in brain metastasis samples vs. primary and ECM samples, using a previously validated Gaussian process model ^22^. Homozygous deletions of 9p21.3 and 2q35, as well as amplification of 12q15 were under positive selection in RCC brain metastases compared extracranial metastases and primary tumors (**Fig. 1E**).

We also looked for recurrent single nucleotide variants (SNVs) in RCC ^16^. We identified *VHL, PBRM1, TP53, MTOR, SETD2* genes as significantly frequently mutated in all three groups. However, the mutation frequencies in the genes were not significantly different among RCC primary tumors, extracranial metastases, or brain metastases (**Supplementary Fig. 1D**).

### Matched-pair analyses of RCC brain metastases supported candidate CNAs

To further evaluate the candidate genomic alterations associated with brain metastatis, we compared brain metastases samples against the matched primary or extracranial tumor samples (**Fig 1A**). Shared and compartment-specific genomic alterations were identified in these matched pairs (**Fig 2A**). Mutations in well-established RCC genes, including *VHL*, *PBRM1*, *BAP1*, *MTOR*, *TP53*, and *KDM5C*, were shared by brain metastasis and matched ECM/primary samples (**Fig. 2A, 2D**). Matched-pair analyses on the CNA events revealed that gains of 1q, 7q, 7p, and 8q, as well as losses of 4q, 10q, and 14q were significantly enriched in brain metastases compared against matched primaries or extracranial metastases (**Fig. 2B-C**). Indeed, gains of chromosome 7 and 8q, as well as losses in 4q and 14q were significantly enriched under both case-control and matched-pair analyses; these results highlight these events as potential CNAs associated with brain metastasis in RCC (**Supplementary Table 2**).

**Figure 2.**
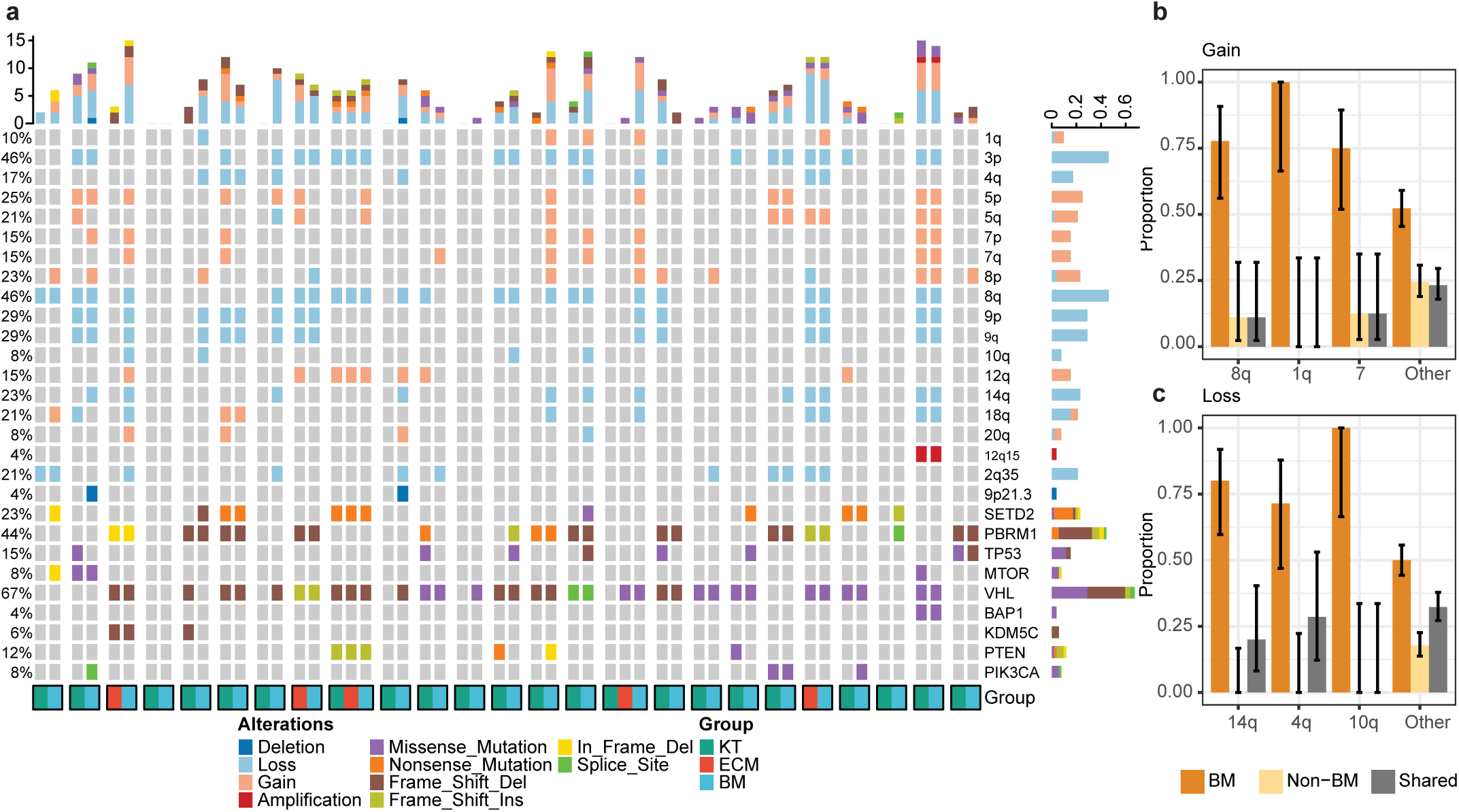
A. Oncoplot of genetic alterations across samples are displayed. Frequencies of alterations for each gene or region are shown to the right. **B-C**. Frequency of significant chromosomal arm level-copy number gains (**B**) and losses (**C**) that are private to the primary or ECM tumor, private to brain metastasis or shared. Proportion of patients with the alterations showing the significance based on Mid-p McNemar test. Binomial confidence interval 1q is present in 4/25 brain metastasis (∼16% abundance in brain metastases) and 0/25 primary/ECM samples. 100% of 1q gain is unique in brain metastases. Similarly, 75% of all 8q gain is unique in brain metastases. Mid-p McNemar test compares matched samples proportion alterations. When any of the samples do not have alteration test filters them out and compares only patient with alteration including their matched samples. FDR< 10%.

### Co-occurrence patterns of genomic alterations in RCC brain metastases

We next characterized the co-occurrence pattern of the candidate CNAs associated with brain metastasis. Several events including 8p loss, 14q loss, 7p gain, and 8q gain demonstrated a strong co-occurrence with each other (**Fig. 3A**). Significant pairwise co-occurrences were observed between 8p loss and 14q loss, 8p loss and 8q gain, as well as 7p gain and 7q gain. From these associations, we constructed a co-occurrence network plot (**Fig. 3B**), which identifies 8p loss as a key hub, suggesting its pivotal role in the development of brain metastasis.

**Figure 3.**
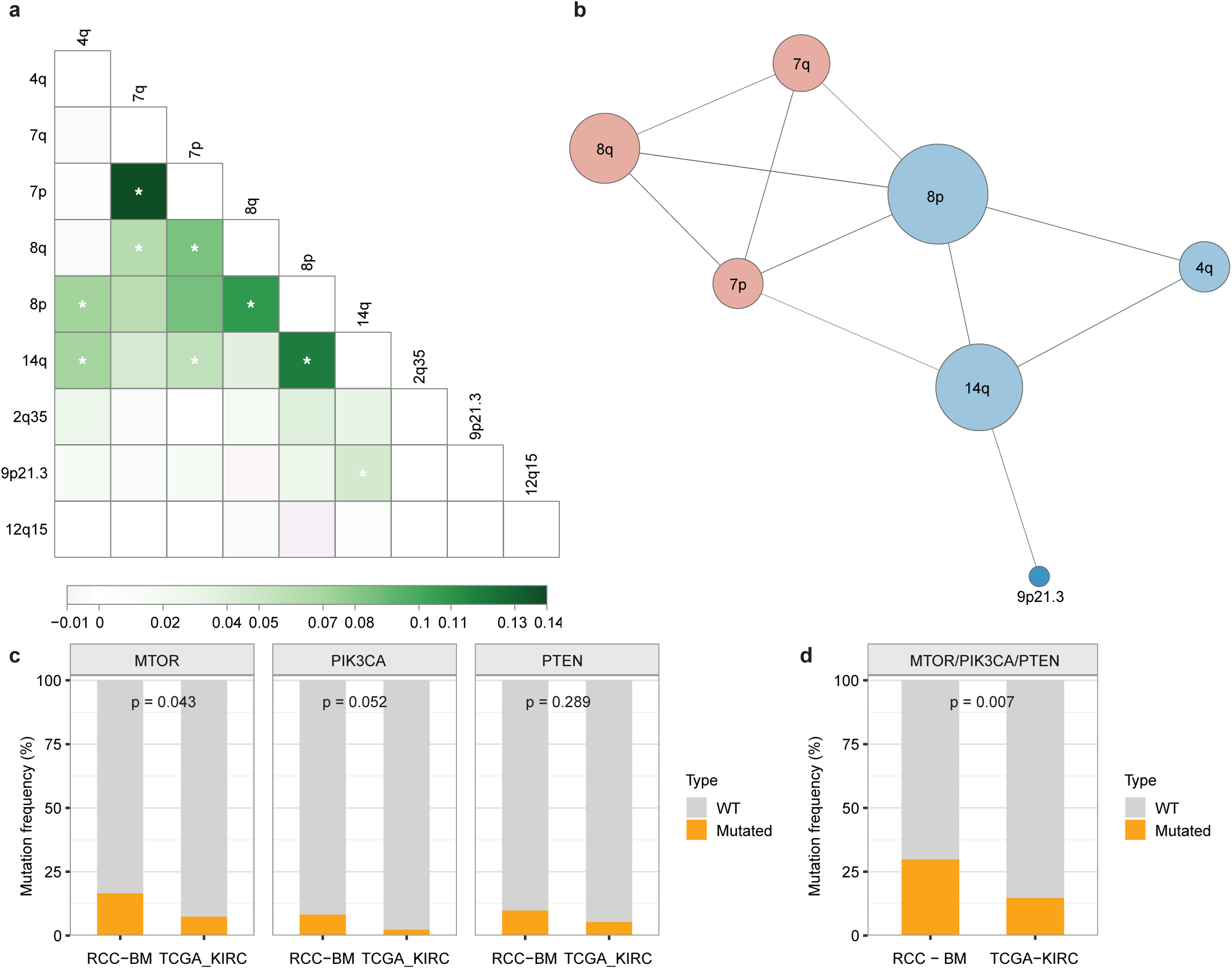
**A**. Pairwise co-occurrence heatmap of copy number alterations (CNAs) across chromosomal regions in the dataset. The x- and y-axes represent chromosomal regions, with color intensity indicating the degree of co-occurrence between alterations. Significant co-occurrence relationships are marked by stars (*). Positive co-occurrence effect sizes are shaded green, while negative correlations are shaded white or gray. **B**. Network plot depicting relationships among frequently altered chromosomal regions. Each node corresponds to a chromosomal region, with red nodes representing gains and blue nodes indicating losses. Node size reflects the frequency of alterations, and edges represent significant co-occurrence or mutual exclusivity relationships. Line thickness between nodes is proportional to the p-value, highlighting the strength of the association. **C**. Bar plots showing mutation frequencies for MTOR, PIK3CA, and PTEN in RCC-BM (Renal Cell Carcinoma - Brain Metastasis) and TCGA-KIRC (The Cancer Genome Atlas - Kidney Renal Clear Cell Carcinoma). Wild-type (WT) cases are shown in gray, and mutated cases in orange. P-values indicate the statistical significance of differences in mutation frequencies between the two groups. **D**. Bar plot illustrating the combined mutation frequency of the MTOR/PIK3CA/PTEN pathway in RCC-BM and TCGA-KIRC groups. Orange bars represent samples with mutations in at least one gene of the pathway, while gray bars represent wild-type cases. The associated p-value reflects the significance of the differences between the groups.

To determine whether the co-occurrence patterns observed in brain metastases are already detectable in primary tumors, we examined pairwise CNA co-occurrence in TCGA-KIRC primary RCC (**Supplementary Fig. 2**). The analysis revealed several significant co-occurring events, consistent with findings from our RCC BM cohort. Notably, the loss of 9p21.3 frequently co-occurred with the loss of 14q (P < 0.0001), while the gain of 8q frequently co-occurred with loss of 8p. These co-occurrence patterns of events in primary tumors suggest that these genomic constellations are established early and may predispose to subsequent brain metastatic dissemination.

Arm-level CNAs may cooperate with pathway-activating point mutations to drive brain metastasis. The PI3K/AKT/mTOR pathway has been recurrently implicated in brain metastasis across breast cancer, lung cancer, and melanoma, and several candidate CNA regions (e.g., PTEN on 10q) overlap with this pathway ^8,23–27^. We therefore looked at whether PI3K/AKT/mTOR pathway mutations were enriched in RCC brain metastases compared to primary tumors. Analysis of our RCC brain metastases revealed higher combined mutation frequencies of *MTOR*, *PIK3CA*, and *PTEN* compared to the primary RCC tumors (P = 0.007; **Fig. 3C-D**).

### Recurrent CNAs associated with brain metastasis are linked with poor patient survival

To assess the prognostic significance of the candidate CNAs enriched in brain metastasis in RCC (2q25 loss, 4q loss, 7p gain, 7q gain, 8p loss, 8q gain, 9p21.3 loss, 12q15 gain, and 14q loss), we conducted survival analyses using the TCGA-KIRC cohort. We first identified 147 patients whose tumors lacked all of the candidate CNAs, and we used this group as the reference in the survival analyses (**Fig. 4A**). Except for 2q35 loss, the candidate CNAs were associated with poor prognosis (**Fig. 4B**, **Supplementary Fig. 3A**). Collectively, patients who presented with one or more of the prognostically significant alterations had poorer overall survival (P=0.004) compared to the reference (**Fig. 4D**). We next looked at the impact of the candidate CNAs on progression-free interval (PFI). Similarly, all CNAsexcept for 2q35 loss were associated with shorter PFI (**Fig. 4C**). Patients with RCC tumors harboring one or more of the alterations exhibited significantly shorter PFIs (P < 0.0001; **Fig. 4E**). Given that most samples without candidate alterations were stage I tumors, we performed further survival analyses on stage I patients. This analysis showed a trend toward poorer overall survival among Stage I RCC patients with the candidate CNAs, although the difference did not reach statistical significance (P=0.06), likely due to the limited number of death events observed among Stage 1 patients (**Supplementary Fig. 3A-B**).

**Figure 4.**
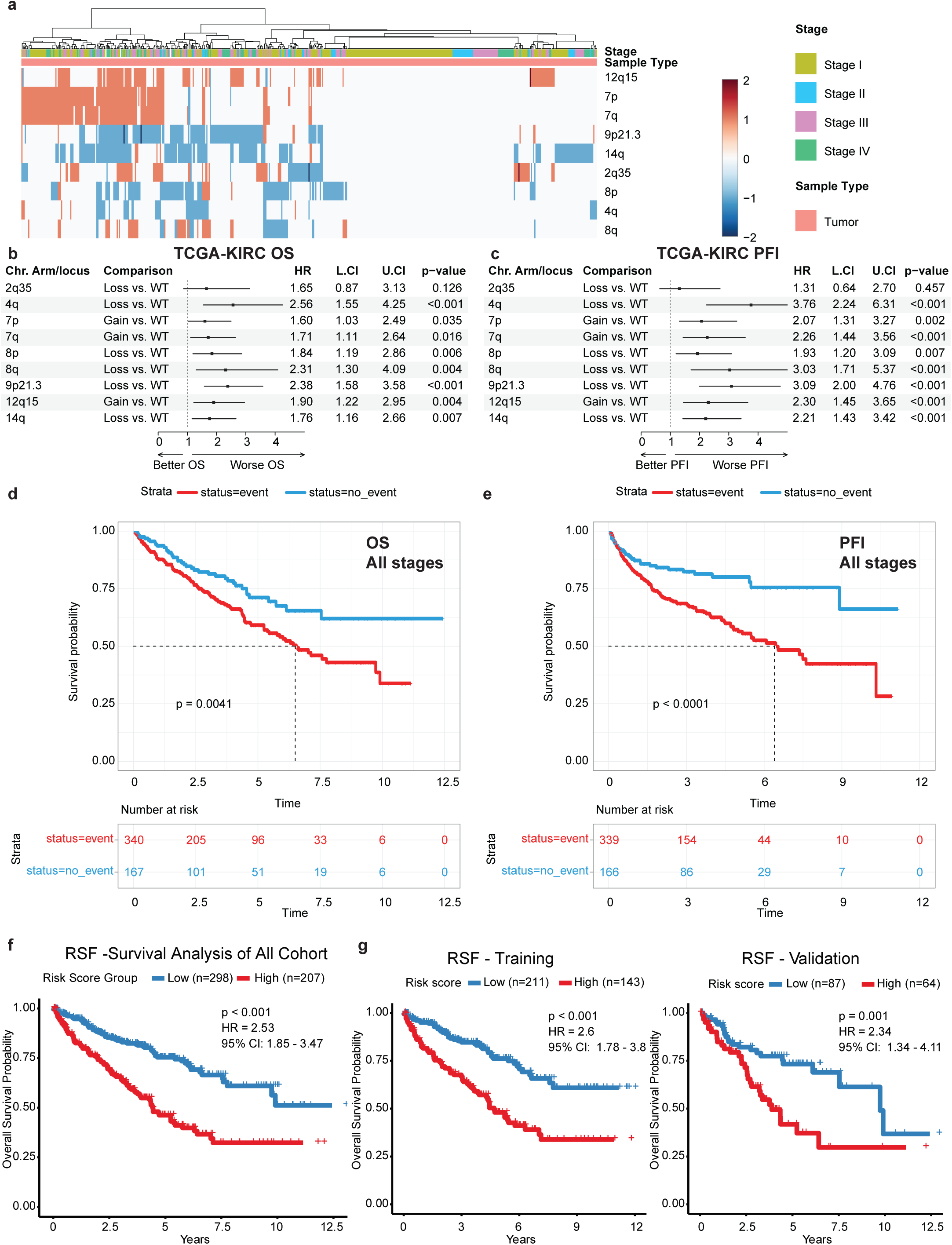
**A**. Heatmap shows CNAs within the TCGA-KIRC cohort, categorized by stage (Stage I-IV). Gains and losses are shown in red and blue, respectively, with intensity indicating the degree of alteration. **B-C**. Forest plots showing the hazard ratios (HR) for overall survival (OS) (**B**) and progression-free interval (PFI) (**C**) associated with chromosomal arm alterations. **D-E.** Kaplan-Meier survival curves show the impact of having one or more significant CNAs on OS (**D**) and PFI (**E**) in patients across all stages. The red survival curve represents patients with events (alterations), while the blue curve represents those without. (**F**) Prognostic performance of the RSF model based on CNV events in TCGA-KIRC. Kaplan–Meier survival analysis of the combined cohort (training and validation) showing significantly poorer overall survival in the high-risk group (HR = 2.53, 95% CI: 1.85–3.47, p < 0.001). (**G**) Separate analyses of the training and validation cohorts demonstrated consistent prognostic separation (Training: HR = 2.60, 95% CI: 1.78–3.81, p < 0.001; Validation: HR = 2.34, 95% CI: 1.34–4.11, p = 0.001).

Additionally, we developed a random survival forest model based on the CNV events (4q, 7p, 7q, 8p, 8q, 9p21.3, 12q15, and 14q), and we stratified TCGA-KIRC patients into prognostically distinct groups. Survival analysis revealed significantly poorer overall survival in the high-risk group than in the low-risk group (HR = 2.53, 95% CI: 1.85–3.47, p < 0.001; **Fig. 4F**). To assess the potential of model overfitting, we randomly split the patients into training and validation sets. Indeed, consistent survival differences were observed in the training (HR = 2.60, 95% CI: 1.78–3.81, p < 0.001) and validation sets (HR = 2.34, 95% CI: 1.34–4.11, p = 0.001) (**Fig. 4G**). The model yielded a mean concordance index (C-index) of 0.64 across the training and validation sets.

### Functional evidence for the role of 8p loss, 8q gain, and 9p21.3 deletion in promoting brain metastasis

No RCC-specific experimental metastasis mapping dataset currently exists. We therefore leveraged MetMap500 data from lung adenocarcinoma (LUAD), breast cancer (BRCA), and melanoma to test whether the same CNAs identified in RCC brain metastases are associated with brain-tropic metastatic potential across cancer types (**Fig. 5A**). In the MetMap study, investigators injected a pool of 500 barcoded cancer cell lines intracardially into mice to allow spontaneous metastasis to distant organs, and harvested organs were analyzed by RNA sequencing to quantify barcode abundance. By integrating MetMap with copy-number profiles of the cell lines from DepMap, we identified genomic alterations that are associated with the potential of the cell lines to metastasize to the brain and other organs ^17^.

**Figure 5.**
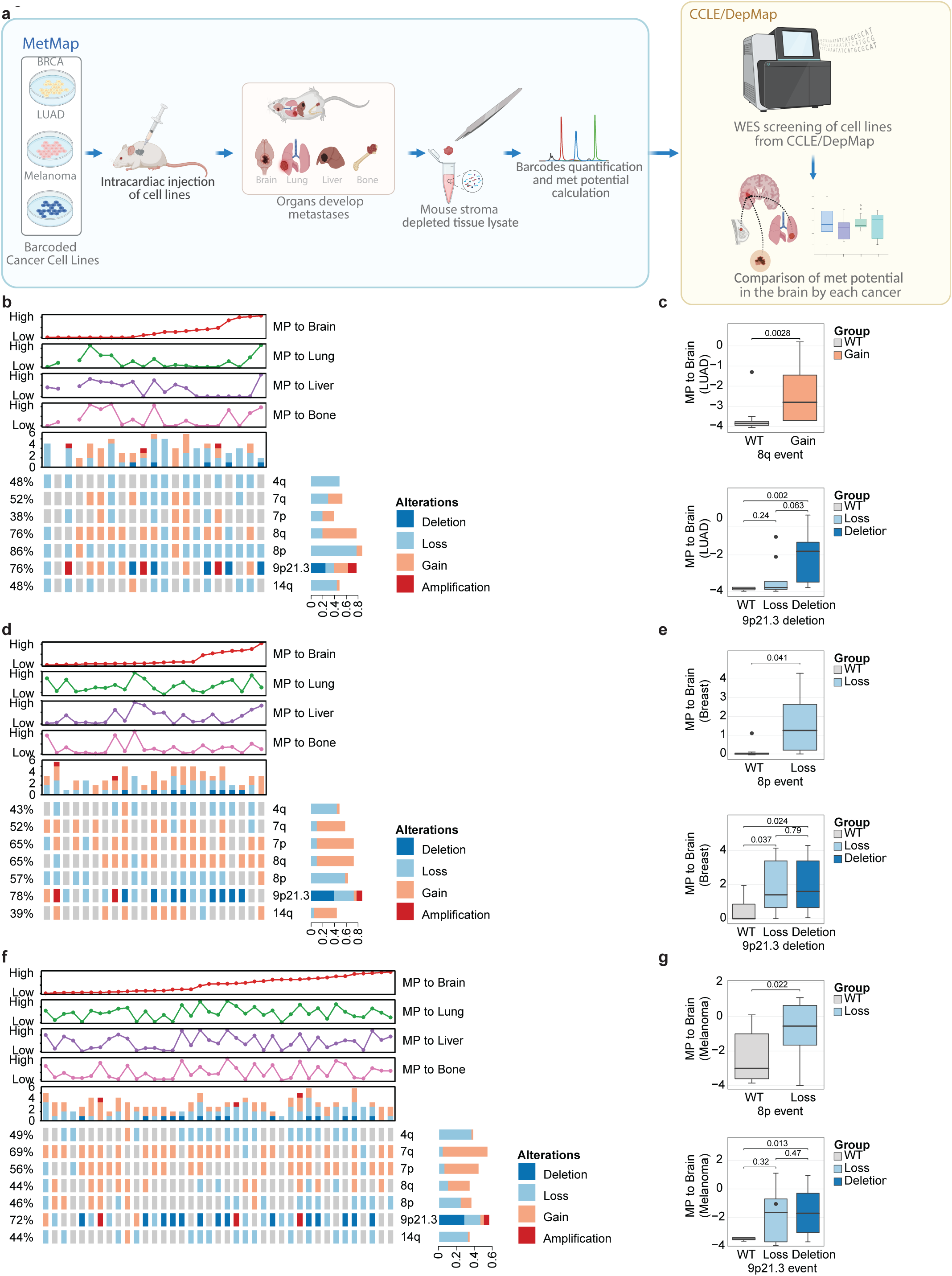
A. An overview of the experimental approach and findings regarding organotropic metastases in Lung Adenocarcinoma (LUAD), Breast Cancer (BRCA), and melanoma. The figure outlines the methodology, starting with intracardiac injection of barcoded cancer cell lines into mouse models, followed by the collection of metastatic tissues from the brain, lung, liver, and bone. DNA was extracted from these tissues, and whole-exome sequencing (WES) was performed to identify genomic alterations. The data were analyzed to compare metastasis-prone (MP) subclones with the parental cell line, revealing potential drivers of metastasis. Genomic alterations in metastatic samples from LUAD (**B**), BRCA (**D**) and melanoma (**F**) cell lines are shown across different organ sites, including the brain, lung, liver, and bone. Line graphs display the distribution of alterations categorized by metastatic potential (high, average, or low). Heatmaps illustrate the distribution and frequency of the candidate driver chromosomal alterations identified in RCC BM. Statistically significant chromosomal alterations associated with MP to the brain are highlighted for LUAD (**C**), BRCA (**E**), and melanoma (**G**) cell lines.

We found that LUAD cell lines with higher brain metastatic potential predominantly exhibit gains in 8q and deletions in 9p21.3 (**Fig. 5B-C**). In contrast, compared against wildtype, these alterations were not associated increased metastatic potentials to the liver, bone, or lung (**Supplementary Fig. 4A, D**). Similarly, BRCA cell lines with 8p loss and 9p21.3 loss demonstrated significantly higher brain metastatic potential compared to WT cells. (**Fig. 5D-E**). There was also no significant differences in the lung, liver, bone metastasis potentials of 8p and 9p21.3 loss cell lines compared to wildtype (**Supplementary Fig. 4B, E**). In melanoma cell lines, we observed a similar pattern (**Fig. 5F**), where both loss of 8p and the deletion of 9p21.3 were significantly associated with increased metastatic potential to the brain, but not to the liver, bone, or lung (**Fig. 5G**, **Supplementary Fig. 4E-F**).

Overall, our results suggest two sets of genomic alterations that are associated with brain metastases: 8q gain with 9p21.3 deletion, and 8p loss with 9p21.3 deletion. Interestingly, the curves for metastatic potential to the liver, lung, and bone generally follow a similar pattern across the three cancer types, while the metastatic potential to the brain seems to follow a different trend. Taken together, our results suggest that 8q gain, 8q loss, and 9p21.3 deletion can enhance metastatic potential to the brain across different cancer types.

### Pan-cancer analysis confirmed the roles of 8p loss, 8q gain, and 9p21.3 deletion in brain metastases across different cancers

To externally validate the candidate CNAs (8p loss, 8q gain, 9p21 loss) associated with brain metastasis, we conducted a comprehensive analysis across cancer types from MSKCC cohorts ^19^ (**Supplementary Fig. 5A**). Samples included in our discovery cohort for case-control analyses were removed from this external validation cohort. Significantly enriched cytogenetic alterations were identified using weighted logistic regression with matching weights, which were determined by coarsened exact matching to ensure exact balance of confounding covariates, including tumour type, as previously described ^11^. Brain metastases (n=475) exhibited significantly higher frequencies of 8p loss, 9p21.3 deletion, and 8q gain, compared to primary tumors (n=15,030) and extracranial metastases (n=10,089) (**Fig. 6A**). These findings suggest a critical role of these alterations in promoting brain metastases. A distinct cluster of brain metastases is characterized by concurrent 9p21.3 deletion and chr8 alterations (**Fig. 6B**).

**Figure 6.**
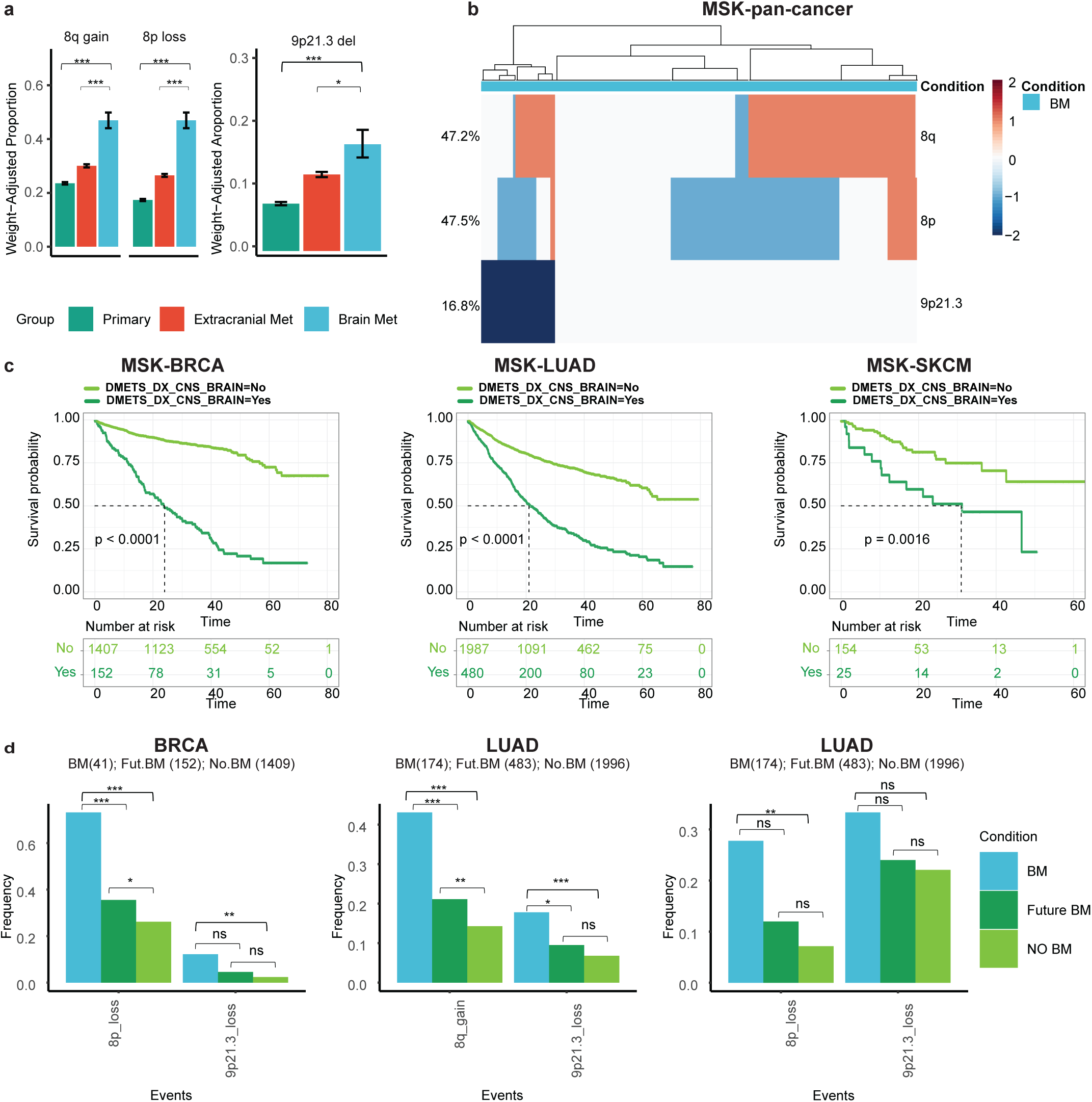
A. Weight-adjusted proportions of 8q gain, 8p loss and 9p21.3 deletion across primary tumors, extracranial metastases, and brain metastases in pan-cancer MSK cohort. Significantly enriched cytogenetic alterations were determined using weighted logistic regression using matching weights. **B.** A hierarchical clustering heatmap demonstrates the frequency of genomic alterations across brain metastases. Alterations include losses (blue) and gains (red) for regions such as 8q, 8p, and 9p21.3. Clustering indicates patterns of co-occurrence among alterations in metastatic samples. **C.** Survival analyses demonstrate the impact of specific genomic alterations on overall survival (OS) in breast cancer (BRCA), lung adenocarcinoma (LUAD and melanoma (SKCM). Patients with the alterations (light green curve) are compared to those without (dark green curve), with significant p-values (e.g., p < 0.001) denoting worse survival outcomes for altered groups. **D.** Bar plots illustrate the frequency of genomic events (8q gain, 8p loss, 9p21.3 loss) in the three cancer types in different groups: in brain metastasis samples (BM), primary tumors from patients who would develop brain metastasis in the future (future BM), and primary tumors from patients who never developed brain metastasis (No BM). Statistically significant differences between conditions after Chi Square test are marked by asterisks (*). Forest plots show the hazard ratios (HR) for overall survival associated with specific genomic alterations for breast cancer (MSK-BRCA), lung adenocarcinoma (MSK-LUAD) and melanoma (MSK-SKCM) in MSK cohort

We analyzed the survival of patients with brain metastases in BRCA, LUAD, and melanoma in MSK MetTropism dataset. Patients with brain metastasis had worse overall survival consistently across cancer types (**Fig. 6C, Supplementary fig. 5B**), highlighting the prognostic significance of brain metastases.

We next examined the frequencies of the candidate CNAs in brain metastasis samples (BM), primary tumors from patients who would develop brain metastasis (Future BM), and primary tumors from patients who never developed brain metastasis (No BM) in breast cancer, lung cancer, and melanoma during the follow-up period (30 months) ^19^. Indeed, the frequencies of 8p loss, 8q gain, and 9p21 deletion were consistently highest in the BM group (**Fig. 6D**). Furthermore, the Future BM group showed intermediate frequencies these alterations, suggesting that these alterations may serve as early indicators of potential brain metastasis. We observed similar findings in colorectal cancer (MSK-COAD), small-cell lung cancer (MSK-SCLC), and prostate cancer (MSK-PRAD) (**Supplementary Fig. 5C**).

We next performed survival analyses to assess the prognostic significance of 8p loss, 8q gain, and 9p21.3 deletion across cancer types (**Supplementary Fig. 6A-B**). In breast cancer, 8p loss was significantly associated with worse OS, with (HR: 1.55, 95% CI: 1.23-1.95, P < 0.001), and 9p loss similarly showed a significantly worse prognosis (HR: 1.79, 95% CI: 1.40-2.29, P < 0.001). In lung cancer, 8q gain, 9p21.3 deletion, and 9p loss were significantly associated with worse OS, with hazard ratios of 1.57 (HR: 1.57, 95% CI: 1.33-1.87, P < 0.001), 1.83 (95% CI: 1.46-2.30, P < 0.001) and 1.63 (95% CI: 1.38-1.92, P < 0.001), respectively. In melanoma, both 9p21.3 deletion and 9p loss were significantly associated with worse OS with hazard ratios of 2.19 (95% CI: 1.39-3.46, P < 0.001) and 1.83 (95% CI: 1.21-2.76, p = 0.004), respectively.

Although 8p loss showed a trend toward worse OS, it was not statistically significant (HR: 1.41, 95% CI: 0.85-2.37, p = 0.187).

To further validate the candidate CNAs enriched in brain metastasis, we also conducted the same survival analyses in colorectal cancer (MSK-COAD), small-cell lung cancer (MSK-SCLC), and prostate cancer (MSK-PRAD). The results were largely consistent with our findings for the other cancer types above (**Supplementary Fig. 7A-B**). Above all, 8p loss, 8q gain, and 9p21.3 loss are consistently associated with brain metastasis and poor prognosis multiple cancer types.

## DISCUSSION

We identified several candidate genomic alterations associated with brain metastases in RCC, including 8p loss, 8q gain, and 9p21.3 deletion. Beyond RCC, the candidate CNAs also occur frequently in brain metastases from lung adenocarcinoma, breast cancer, and melanoma, potentially indicating shared mechanisms of brain metastatic dissemination across cancer types. Importantly, these genomic events were associated with poor prognosis even in stage I RCC, highlighting their potential value for early risk stratification and personalized monitoring.

Our results, along with findings in other studies, collectively suggest that 9p21.3 deletion contributes to the formation of brain metastases from RCC and other cancer types. This is important because 9p21.3 deletion occurs frequently in ∼13% of cancers ^28^. The 9p21.3 locus contains *CDKN2A/B* and *MTAP*, key regulators of cell cycle control and tumor metabolism ^29,30^. Alterations in *CDKN2A/B* have been implicated in brain metastases across NSCLC ^31^, LUAD ^11^, breast cancer ^32^, and melanoma ^33^, and CDKN2A/B co-deletion in resected brain metastases correlates with increased local and distant CNS progression ^34^. Recent work shows that 9p21.3 deletion promotes a “cold” immune microenvironment through type I IFN pathway suppression and loss of regulatory elements ^28,35^, contributing to poor responses to immune checkpoint therapy ^28,35^. Notably, RCC tumors with 9p21 deletions exhibit CD8+ T-cell infiltration yet still experience poor survival after PD-1 blockade ^36^, potentially explaining the limited immunotherapy benefit in RCC brain metastasis ^37^ and challenging the assumption that immune infiltration predicts improved outcomes ^38^. Functionally, 9p21.3 loss appears to be an early metastasis-enabling event. In pancreatic cancer, loss of type I IFN genes in this region promotes immune evasion during metastatic seeding ^39^, and 9p21 loss has also been shown to confer a chromosomal-instability–tolerant phenotype that supports aneuploid tumor growth and metastasis ^40^. Together, these findings support 9p21.3 deletion as an early genomic trigger in metastatic progression, especially to the brain. Therapeutically, CDK4/6 inhibitors are being evaluated to target CDKN2A/B loss in brain metastases (NCT02896335; NCT03994796).

Gain of 8q is also a promising driver of RCC brain metastasis. It is associated with metastasis and poor survival in ccRCC ^41^. Multiple studies have shown focal amplification of 8q24 in ∼15–23% of ccRCC, consistently involving *MYC* within the amplified region ^18,42,43^. MYC amplification has previously been identified and functionally validated as a brain metastatic driver in lung adenocarcinoma ^11^.. Mechanistically, MYC promotes brain colonization by enhancing invasiveness, recruiting macrophages, and inducing gap junction formation between cancer cells and astrocytes ^44^. MYC also upregulates redox enzymes that mitigate ROS generated by activated microglia, potentially protecting metastatic cells within the brain microenvironment ^45^. Several preclinical studies and early-phase clinical trials are currently investigating MYC-targeting approaches, but none are specifically designed for patients with brain metastases ^46^.

We also observed frequent 8p loss in RCC brain metastases, which has been associated with metastatic potential in multiple cancer types ^47,48^. Evidence suggests that 8p loss promotes metastasis through deregulation of multiple tumor suppressors including MSRA, NAT1, PPP2CB, and DLC1, rather than a single driver ^48–50^. In prostate cancer, loss of NKX3.1 (also on 8p) increases oxidative stress and disrupts mitochondrial function, thereby promoting progression ^51^. Notably, the 8p-loss cluster was enriched in breast cancers that developed brain, but not lung or bone, metastases ^17^. It alters fatty acid and ceramide metabolism, increasing invasiveness and autophagy ^49^. Together, these findings suggest that 8p loss may contribute to biological processes relevant to brain metastasis, although causal mechanisms remain to be fully elucidated.

We observed significant associations of these recurrent CNAs in brain metastasis across multiple cancer types, supporting the concept of shared genomic determinants of organ-specific metastasis. This is consistent with the pan-cancer brain metastasis genomic analysis by Brastianos et al.^8^, which identified recurrent PI3K/AKT pathway alterations across brain metastases from diverse primary sites. However, whether these alterations directly confer brain tropism or reflect broader features of aggressive disease remains to be determined.

To translate these genomic insights into a clinically relevant framework, we developed a RSF model incorporating recurrent CNAs.). To our knowledge, this is the first CNA-based prognostic model in RCC, achieving a concordance index of 0.64, comparable to established clinical models such as IMDC, highlighting the additive prognostic value of genomic data ^52^. Therefore, our CNA-based survival model is promising candidate for future validation in prospective cohorts.

This study has some limitations. First, copy-number and SNV calls were derived from bulk WES, which cannot fully resolve intratumoral heterogeneity or spatially restricted clones within the brain microenvironment. Second, functional validation is indirect: MetMap analyses rely on cell-line systems and intracardiac injection models that may not recapitulate all aspects of human CNS colonization, including immune contexture, blood–brain barrier dynamics, and stromal–neural interactions. Third, RCC-specific experimental models of brain metastasis remain limited, restricting direct mechanistic validation of the observed associations.

In summary, we identify recurrent CNAs that are enriched in RCC brain metastasis and associated with adverse clinical outcomes across multiple cancer types. These findings suggest a potential genomic basis underlying brain metastasis and support further investigation of CNA-based biomarkers for risk stratification. Our results uncover new areas of brain metastasis research, such as dissecting how 8p loss, 8q gain, and 9p21.3 deletion cooperate to promote brain tropism by using CNS-competent PDX systems and in vivo CRISPR perturbations ^53^. Another future direction is explore how the CNA drivers remodel tumour microenvironment remodeling. Finally, potential avenues for therapeutic development for brain metastases include CDK4/6 inhibition in *CDKN2A/B*-deleted tumors, synthetic-lethality in MTAP co-deleted disease, and targeting lipid metabolism in RCC with 8p loss.

### List of abbreviations

Brain metastasis (BM); breast cancer (BRCA); central nervous system (CNS); clear cell renal cell carcinoma (ccRCC); concordance index (C-index); copy-number alteration (CNA); extracranial metastasis (ECM); formalin-fixed paraffin-embedded (FFPE); Genome Analysis Toolkit (GATK); hazard ratio (HR); interferon (IFN); International Metastatic Renal Cell Carcinoma Database Consortium (IMDC); institutional review board (IRB); kidney renal clear cell carcinoma (KIRC); lung adenocarcinoma (LUAD); Metastasis Map of Human Cancer Cell Lines (MetMap); Memorial Sloan Kettering (MSK) and Memorial Sloan Kettering Cancer Center (MSKCC); methylthioadenosine phosphorylase (MTAP); non–small cell lung cancer (NSCLC); overall survival (OS); programmed cell death protein 1 (PD-1); progression-free interval (PFI); progression-free survival (PFS); phosphoinositide 3-kinase (PI3K); phosphatase and tensin homolog (PTEN); quantitative polymerase chain reaction (qPCR); reactive oxygen species (ROS); renal cell carcinoma (RCC); renal cell carcinoma brain metastasis (RCC-BM); random survival forest (RSF); single-nucleotide variant (SNV); small-cell lung cancer (SCLC); The Cancer Genome Atlas (TCGA), including the TCGA kidney renal clear cell carcinoma cohort (TCGA-KIRC); wild type (WT); whole-exome sequencing (WES)

## DECLARATIONS

### Author Contributions

Conceptualization: EH

Tissue collection and Data curation: MH, SH, BB, BB, LMN, MM, FG, JTH, AOO, MAB, EJ Formal Analysis: BGK, JAO, ALF, PL, JWTT, DJHS

Funding acquisition: EH, MH

Sample preparation and sequencing: HDC, EJT Project administration: EH

Supervision: EH, DJHS

Visualization: BGK, JAO, ALF, PL, JWTT, DJHS Writing – original draft: BGK

Writing – review & editing: all authors

### Data and Code Availability

Processed DNA sequencing data, and supporting files will be made available through a project-specific request to the corresponding author. The scripts used for the analysis are available at the following GitHub repositories: https://github.com/hasanovlab (available upon publication).

## Acknowledgements

We wish to acknowledge the Pathology Core within the Brain Tumor Research Program at MD Anderson Cancer Center supported through the SPORE in Brain Cancer award (P50CA127001) and the generous philanthropic contributions to the MD Anderson Cancer Neuroscience Program.

## Declaration of generative AI and AI-assisted technologies in the manuscript preparation process

During the preparation of this work the author(s) used ChatGPT in order to improve the grammar and readability of the manuscript. After using this tool/service, the author(s) reviewed and edited the content as needed and take(s) full responsibility for the content of the published article.

